# K_Ca_3.1 Drives Pro-Fibrotic Activation and Represents a Novel Therapeutic Target in Aortic Stenosis

**DOI:** 10.64898/2026.04.30.720379

**Authors:** Molly Whitfield, Saadia Aslam, João Gonçalves de Sousa, Daniel Taveira, Catherine McMullan, Mathuscha Ratnasingham, Gill Elliott, S Mark Duffy, Neil Craig, Stefan Veizades, Stephanie Sellers, Hiwa Sherzad, Metesh Acharya, Giovanni Mariscalco, Gerry P McCann, Peter Bradding, Anvesha Singh, Katy M Roach

## Abstract

**Introduction:** Aortic stenosis (AS) is characterised by progressive aortic valve (AV) leaflet fibrosis and calcification, yet no medical therapies exist to slow disease progression. AV interstitial cells (VICs) that differentiate into myofibroblasts are central drivers of fibrosis. The Ca^2+^-activated K^+^ channel K_Ca_3.1 promotes pro-fibrotic signalling in several fibrotic diseases, however its role in AS remains unknown.

**Methods:** K_Ca_3.1 protein expression was examined in paraffin embedded tissue by Immunohistochemistry from control and AS valve tissue. VICs were isolated, cultured and phenotypically characterised as myofibroblasts from AV tissue obtained from patients with severe tricuspid AS undergoing surgical AV replacement (n=19). K_Ca_3.1 mRNA and protein expression were assessed by qRT-PCR and immunohistochemistry, and functional channel activity confirmed using patch-clamp electrophysiology. The effects of transforming growth factor-β1 (TGFβ1) stimulation and pharmacological inhibition with the selective K_Ca_3.1 blocker senicapoc were examined.

**Results:** Immunoreactive K_Ca_3.1 channels and smooth muscle actin were detected in both control and AS aortic valve tissue, localised to elongated, nucleated interstitial cells, with significantly higher expression observed in AS tissue compared to control. Isolated VICs exhibited an activated myofibroblast phenotype, expressing THY-1, vimentin, collagen and α-smooth muscle actin (αSMA) (n=9). Myofibroblasts expressed K_Ca_3.1 mRNA and protein and demonstrated functional plasma membrane channels. TGFβ1 stimulation increased K_Ca_3.1, αSMA and collagen type I mRNA expression, while K_Ca_3.1 blockade with senicapoc (100 nM) significantly attenuated TGFβ1-induced αSMA expression, stress fibre formation and collagen gel contraction. Senicapoc had no effect on myofibroblast proliferation or migration.

**Conclusions:** We show for the first time that functional K_Ca_3.1 channels are expressed in human AS tissue and AV myofibroblasts, where they regulate myofibroblast contraction, α-SMA expression, and differentiation, promoting pro-fibrotic activity. These responses are attenuated by the selective K_Ca_3.1 inhibitor senicapoc. Given its established safety in phase 3 clinical trials, K_Ca_3.1 inhibition represents a promising and readily translatable anti-fibrotic therapeutic strategy for AS.

## INTRODUCTION

Aortic stenosis (AS) is the most common valvular heart disease requiring intervention in the developed world[1], with increasing prevalence as the population continues to age[1, 2]. It is characterised by progressive fibrocalcific remodelling of the aortic valve (AV), ultimately causing valve obstruction, left ventricular remodelling and, if untreated, heart failure and death[2]. No pharmacological therapies exist to slow or halt disease progression, and surgical aortic valve replacement or transcatheter aortic valve implantation remain the only definitive treatment. This underscores the urgent need for novel therapeutic strategies.

Mechanical stress and disturbed shear forces compromise valvular endothelial barrier integrity, facilitating lipoprotein infiltration and retention in the subendothelium, which then triggers inflammation, oxidative stress and local activation of resident cells within the valve[3]. A hallmark feature is leaflet fibrosis, driven by the differentiation of valvular interstitial cells (VICs) into activated myofibroblasts[4]. These cells drive pathological extracellular matrix (ECM) deposition, increased contractility, and tissue stiffening. Transforming growth factor beta 1 (TGFβ1) is a central growth factor in this process, inducing α-smooth muscle actin (αSMA) expression and ECM synthesis[5].

Ion channels are established regulators of fibroblast activation and fibrosis in multiple organs, including the AV[6, 7]. The Ca^2+^-activated K^+^ channel K_Ca_3.1 (encoded by KCNN4) is a key myofibroblast channel that maintains the membrane potential required for Ca^2+^ influx, a critical signal for myofibroblast differentiation and activity[8]. K_Ca_3.1 has been implicated in fibroblast proliferation, secretion, and contraction in multiple fibrotic diseases, including pulmonary, renal, and rodent models of myocardial fibrosis[9-11], yet its expression in AV tissue and role in human AV fibrosis, AV myofibroblasts and AS pathogenesis remains unknown.

We hypothesised that K_Ca_3.1 contributes to fibrotic remodelling in AS and represents a novel therapeutic target. Here, we investigated the expression of K_Ca_3.1 in human AV tissue and its function in myofibroblasts isolated from the valves of patients with severe AS. We examined whether pharmacological blockade with senicapoc, a clinically tested selective K_Ca_3.1 channel inhibitor, attenuates key pro-fibrotic features of AV myofibroblast activation.

## METHODS

Complete experimental protocols and reagent information will be made available upon publication in a peer-reviewed journal

## RESULTS

### Baseline Characteristics of the AV donors

The demographic, clinical and echocardiographic characteristics of control and AV donors with severe tricuspid AS are summarised in Table 1. These data indicate that the AS donor cohort is representative of the typical clinical profile of patients with severe AS.

**Table 1.** Clinical and echocardiographic parameters of control and AS valve donors.

| **Characteristic** | **AS** | **Controls** |
| --- | --- | --- |
| Number of subjects | 19 | 7 |
| Male (n, %) | 12 (63%) | 5 (71%) |
| Age (years) | 72 ± 1.4 | 62 ± 4.6 |
| Diabetes mellitus (n, %) | 4 (21%) | 1 (14%) |
| Left Ventricular Ejection Fraction (%) | 47 ± 5 | 30 ± 8 |
| Aortic Valve Mean Gradient (mmHg) | 45.5 ± 2.6 |  |
| Aortic valve area (cm<sup>2</sup>) | 0.77 ± 0.06 |  |
| Previous or concomitant Coronary Artery Bypass Graft (n, %) | 2 (5%) |  |

### K_Ca_3.1 is expressed in AS valve tissue

Immunohistochemical analysis demonstrated clear expression of K_Ca_3.1 in human AV tissue, with positive staining localised predominantly to elongated, nucleated interstitial cells within the control and AS valve leaflets (**Figure 1A**). These K_Ca_3.1-positive cells exhibited morphological characteristics consistent with activated VIC/myofibroblasts, including elongated nuclei and prominent cytoplasmic processes. K_Ca_3.1 immunoreactivity was distributed throughout fibrotic and structurally remodelled regions of AS valves, consistent with areas of active ECM remodelling. Quantitative assessment revealed a marked increase in the proportion of K_Ca_3.1-positive nucleated interstitial cells in AS valves compared with non-diseased control valves (n=7 per group; p<0.0001; **Figure 1B**). This significant upregulation indicates that K_Ca_3.1 expression is strongly associated with the diseased valve state and suggests enrichment within activated myofibroblast populations. In contrast, isotype control sections showed no detectable staining, confirming antibody specificity and the validity of the observed signal.

**Figure 1.**
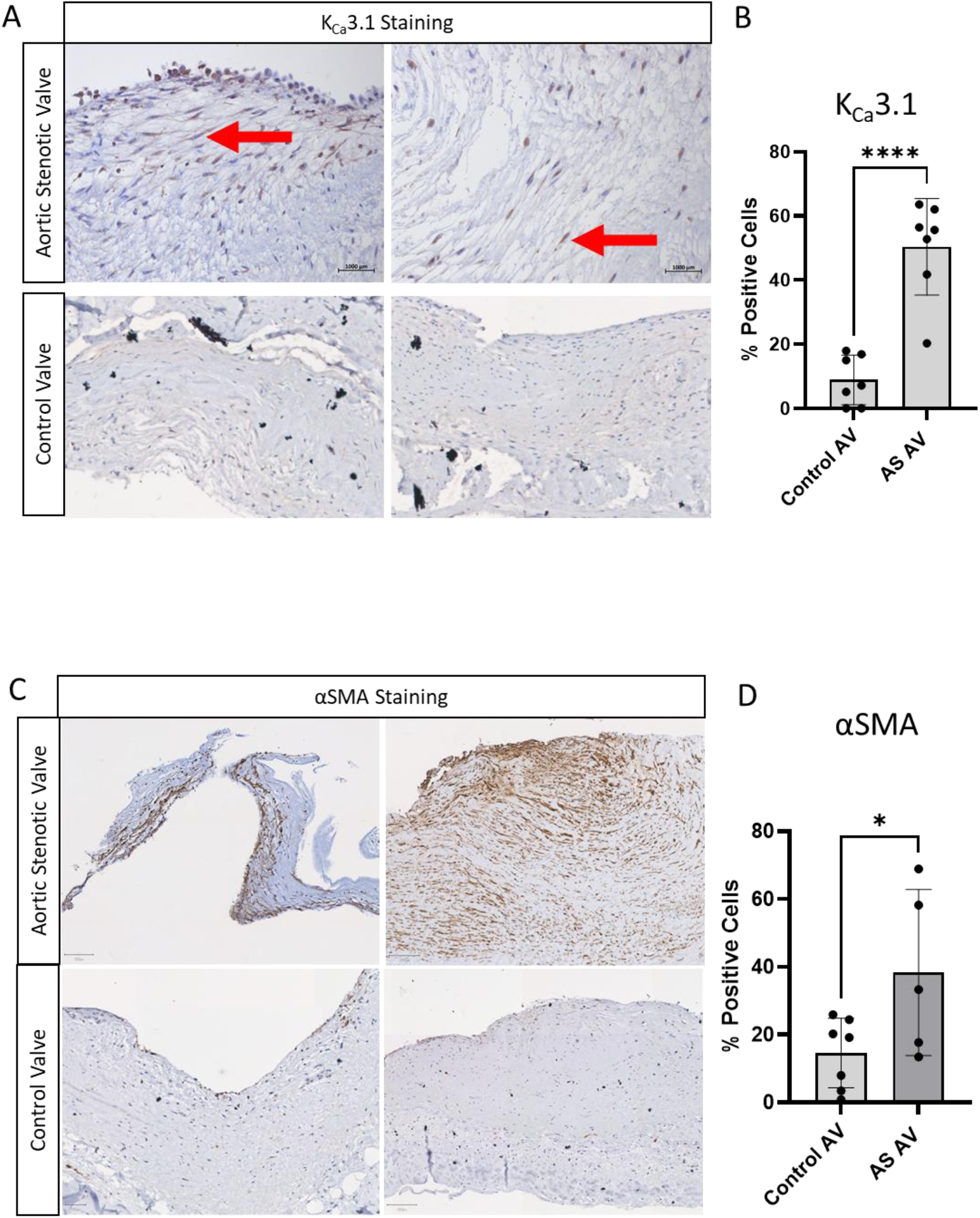
K_Ca_3.1 Expression in Human AS Valve Tissue. A) Representative immunohistochemistry images of AS valve sections showing KCa3.1-positive cells (brown) in multiple donor tissues. Staining was primarily localised to elongated, nucleated interstitial cells distributed throughout the fibrotic valve layers. Red arrows indicate K_Ca_3.1-positive cells with characteristic myofibroblast morphology, including an elongated nucleus and cytoplasmic processes. Sections were counterstained with Gill’s haematoxylin. (B) Quantification of KCa3.1 expression in control and AS valve sections (n=7 control AV, n=7 AS). Bars represent the percentage of K_Ca_3.1-positive nucleated cells per section (mean ± SD), with individual data points shown. ****p<0.0001.(C) Representative immunohistochemistry images showing αSMA-positive cells (brown) in control and AS valve sections. (D) Quantification of αSMA expression in control and AS valve sections (n=7 control AV, n=5 AS). Bars represent the percentage of αSMA-positive nucleated cells per section (mean ± SD), with individual data points shown *p=0.0422.

To further characterise myofibroblast activation, αSMA, a well-established marker of the activated myofibroblast phenotype, was evaluated by immunostaining. Representative sections demonstrated increased αSMA-positive interstitial cells in AS valves compared with controls (**Figure 1C**). Quantitative analysis confirmed a significant increase in the percentage of αSMA-positive nucleated cells in AS valves (control n=7, AS n=5; p=0.0422; **Figure 1D**), indicating enhanced myofibroblast differentiation in diseased tissue.

Collectively, these findings demonstrate that K_Ca_3.1 expression is significantly elevated in AS valves and localises predominantly to cells exhibiting morphological and molecular features consistent with an activated myofibroblast phenotype. The increased abundance of both K_Ca_3.1-positive and αSMA-positive cells in AS valves supports a role for K_Ca_3.1 in myofibroblast activation and valve remodelling associated with disease progression.

### K_Ca_3.1 is expressed by αSMA-positive myofibroblasts in AS valve tissue

Dual immunofluorescence staining demonstrated co-localisation of K_Ca_3.1 with αSMA, a marker of activated myofibroblasts, within interstitial cells of human AS valve tissue (n=4; Figure 2). Whole-section imaging revealed widespread distribution of both K_Ca_3.1 and αSMA throughout the valve leaflets (**Figure 2A,B**). Higher-magnification analysis confirmed that a substantial subset of interstitial cells exhibited overlapping K_Ca_3.1 and αSMA staining, consistent with K_Ca_3.1 expression in activated myofibroblasts (**Figure 2C,D**). Notably, K_Ca_3.1-positive cells were enriched in regions with altered ECM architecture, indicative of fibrotic remodelling (**Figure 2E**). These findings identify K_Ca_3.1 as a feature of myofibroblast-driven tissue remodelling in AS.

**Figure 2.**
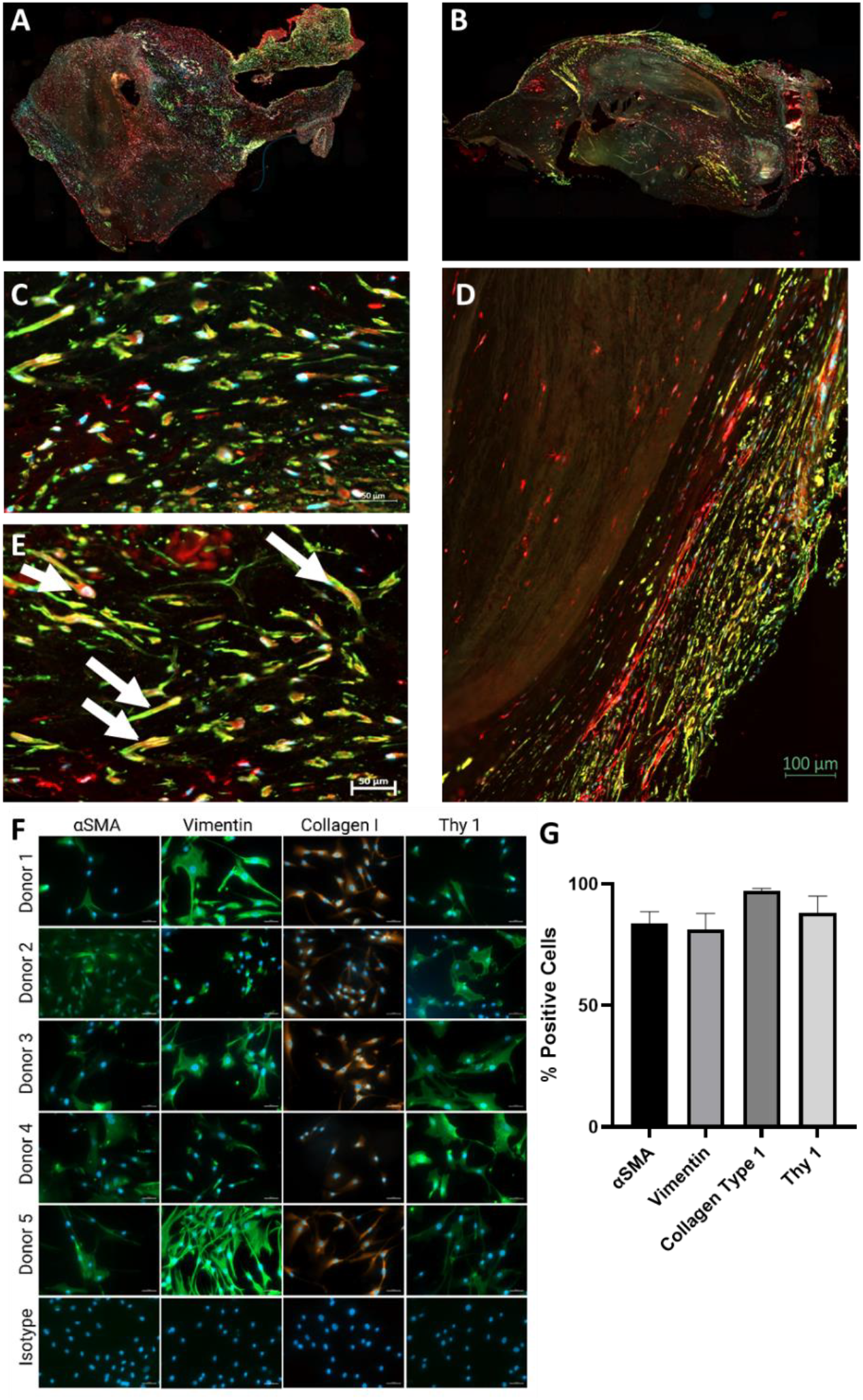
Co-localisation of K_Ca_3.1 with the myofibroblast marker αSMA in AS Valve Tissue. Representative immunofluorescence images of human AV leaflets stained for K_Ca_3.1 (red), αSMA (green), and nuclei (blue) (n=4). (A-B) Whole-section scans showing widespread K_Ca_3.1 and αSMA staining throughout the AV tissue. (C-D) Higher-magnification view of interstitial regions illustrating numerous cells with overlapping K_Ca_3.1 and αSMA staining (yellow), indicating K_Ca_3.1 expression in αSMA-positive myofibroblasts. (E) Lower-magnification image depicting regions of altered ECM within the valve where K_Ca_3.1 positive cells are localised. (F) Representative images from n=5 myofibroblast donors are stained for: α-smooth muscle actin and the isotype control IgG2a; vimentin and isotype control mouse IgG2a; collagen type 1 and rabbit isotype control IgG; and anti-fibroblast antigen (Thy-1/CD90) and the mouse isotype control IgG1. Nuclei were stained with DAPI (blue). Images captured at ×200 magnification. G) Quantification of the percentage of cells expressing αSMA, vimentin, collagen I, and Thy-1 across myofibroblast donors (n=9). Data are presented as mean ± SD.

### Isolation and characterisation of human AV myofibroblasts

Myofibroblasts were successfully isolated from stenotic human AV tissue. Cells displayed characteristic spindle-shaped and stellate morphology consistent with myofibroblasts. Immunofluorescence analysis confirmed expression of established myofibroblast markers (**Figure 2F**). Quantitative analysis across donor isolates demonstrated that 88% of cells expressed the fibroblast marker Thy-1/CD90, 81% expressed vimentin, 97% expressed collagen type I, and 83% expressed αSMA, confirming a predominantly activated myofibroblast phenotype (**Figure 2G**). These findings validate the successful isolation of a highly enriched AV myofibroblast population suitable for functional and mechanistic studies.

### Human AV myofibroblasts express functional K_Ca_3.1 channels that can be pharmacologically modulated

Immunofluorescence staining confirmed robust K_Ca_3.1 protein expression in cultured human AV myofibroblasts (n=6), with punctate staining localised throughout the cytoplasm and consistent with membrane-associated expression, while isotype controls showed no detectable signal, confirming antibody specificity (**Figure 3A**). Consistent with these findings, K_Ca_3.1 mRNA expression was detected in myofibroblasts (n=8) (**Figure 3B**).

**Figure 3.**
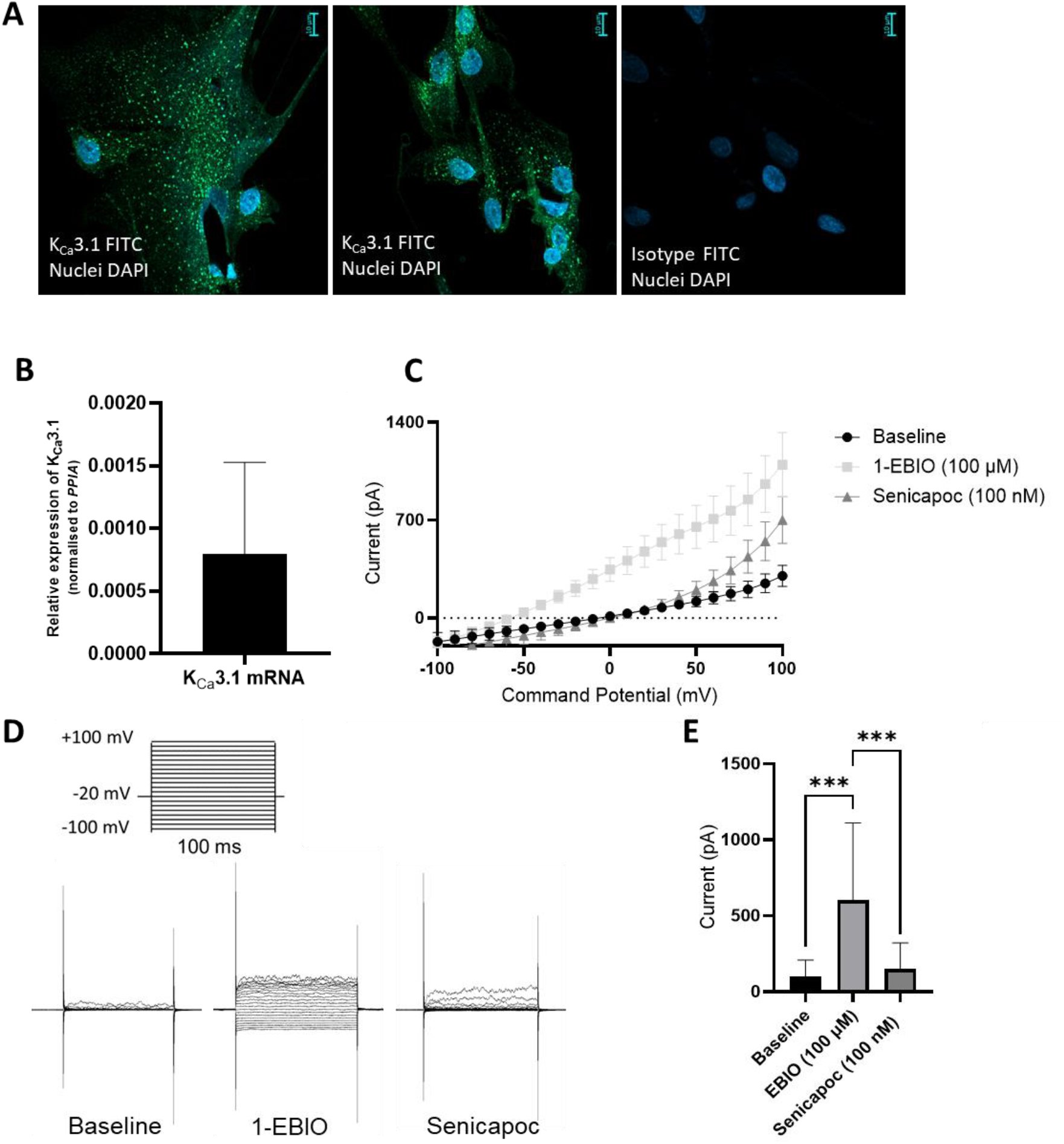
Expression of K_Ca_3.1 in AS valve myofibroblasts. (A) Representative immunofluorescence images showing K_Ca_3.1 protein localisation in cultured human AS valve myofibroblasts (FITC, green); nuclei were counterstained with DAPI (blue). Corresponding isotype control images show no detectable signal, confirming antibody specificity. Images were captured at ×200 magnification; scale bar = 10 µm (n = 6 donors). (B) qRT–PCR analysis of K_Ca_3.1 (KCNN4) mRNA expression in AS valve myofibroblasts, normalised to PPIA. Bars represent mean ± SD (n = 8). C) Current–voltage (I–V) relationship from whole-cell patch-clamp recordings of AS valve myofibroblasts under baseline conditions and following sequential application of the KCa3.1 opener 1-EBIO (100 µM) and the selective blocker senicapoc (ICA-17043, 100 nM) (n = 5 cultures, 13 cells). (D) The voltage protocol and raw current traces from a single cell at baseline, following 1-EBIO, and after senicapoc. (E) Quantification of current amplitude at +40 mV showing a significant increase following 1-EBIO versus baseline, which was significantly reduced by senicapoc. Bars represent mean ± SEM; individual data points are shown (***p=0.0002 for indicated comparisons).

To determine whether K_Ca_3.1 channels were functionally active, whole-cell patch-clamp electrophysiology was performed in isolated AV myofibroblasts. Baseline recordings showed minimal current activity; however, application of the K_Ca_3.1 channel opener 1-EBIO (100 µM) induced robust outwardly rectifying currents across a range of command potentials, with a negative reversal potential, consistent with activation of K_Ca_3.1 channels (**Figure 3C)**. These currents displayed electrophysiological properties characteristic of K_Ca_3.1, with immediate activation with each voltage step and a stable current with no inactivation during a 100 msec pulse.

Importantly, subsequent application of the selective K_Ca_3.1 inhibitor senicapoc (100 nM [IC_50_ 6-10 nM]) markedly reduced 1-EBIO–induced currents, confirming channel specificity (**Figure 3D**). Quantitative analysis demonstrated a significant increase in current amplitude following 1-EBIO stimulation compared with baseline, which was markedly attenuated by senicapoc (p<0.001; **Figure 3E**). These findings confirm that functional K_Ca_3.1 channels are expressed in human AV myofibroblasts.

### K_Ca_3.1 channel inhibition does not affect cell viability

Although oral administration of senicapoc is known to be well tolerated by humans in clinical trials in patients with sickle cell anaemia[12], we sought to determine whether it exhibited any cytotoxic effects in AV myofibroblasts. Cells were cultured in serum-free medium with or without senicapoc (10 or 100 nM), and viability was assessed. As shown in **Figure 4A**, senicapoc treatment at either concentration did not reduce cell viability compared with serum-free controls, indicating that senicapoc is not cytotoxic under these experimental conditions.

**Figure 4.**
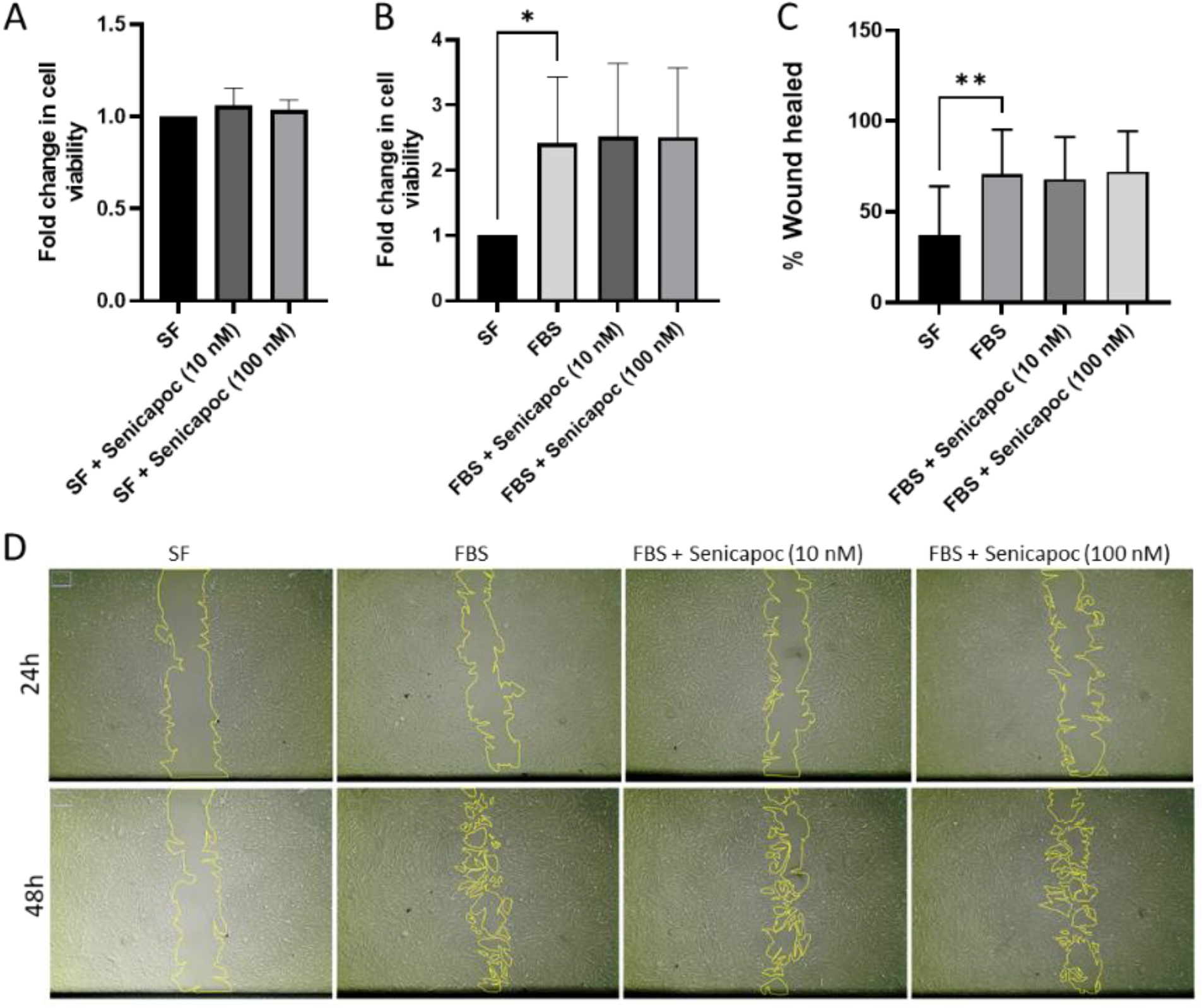
K_Ca_3.1 blockade does not inhibit viability, proliferation, or migration of AV myofibroblasts. (A) Cell viability in serum-free (SF) medium with or without senicapoc (10 or 100 nM). Senicapoc had no cytotoxic effect and the fold change in viability was comparable to the SF control. (B) Proliferation in the presence of FBS ± senicapoc (10 or 100 nM). FBS significantly increased proliferation relative to serum-free (*p<0.05), while senicapoc at either concentration did not further affect the FBS response. (C) 2D wound-healing assay showing percentage of wound closure in serum-free, FBS, and FBS ± senicapoc (10 or 100 nM) for 48h. FBS significantly enhanced wound healing (**p<0.01), senicapoc had no effect. (D) Representative images of wound healing.

### K_Ca_3.1 channel inhibition does not attenuate AV myofibroblast proliferation or migration

K_Ca_3.1 channel inhibition has been implicated in myofibroblast pro-fibrotic function in pulmonary fibrosis[9]. To determine whether K_Ca_3.1 channel inhibition influences key functional properties of AV myofibroblasts, we first assessed the effect of senicapoc on cell proliferation by flow cytometry. As expected, FBS significantly increased cell proliferation relative to serum-free conditions (p<0.05), however, the addition of senicapoc (10 or 100 nM) did not modulate this (**Figure 4B**). We next examined whether K_Ca_3.1 inhibition affected cell migration using a two-dimensional wound-healing assay/scratch assay. FBS markedly enhanced wound closure compared with serum-free medium (p<0.01), but senicapoc at either concentration did not impact the percentage of wound healed over 48h (**Figure 4C and D**). Together, these findings indicate that senicapoc is not cytotoxic and does not modify the proliferative or migratory capacity of AV myofibroblasts under the conditions tested.

### TGFβ1 upregulates K_Ca_3.1 and profibrotic markers in AV myofibroblasts

To determine whether TGFβ1 regulates K_Ca_3.1 expression in AV myofibroblasts, AV myofibroblasts were stimulated with TGFβ1 (10 ng/ml) for 72 h, and transcriptional changes were quantified by qRT-PCR. TGFβ1 significantly increased K_Ca_3.1 mRNA relative to vehicle controls (p<0.05) (**Figure 5A**). TGFβ1 stimulation also led to a robust upregulation of αSMA (p=0.004) (**Figure 5B**) and collagen type I (p=0.005), markers of myofibroblast differentiation (**Figure 5C**). Together, these data demonstrate that TGFβ1 drives increases in K_Ca_3.1 expression and concomitantly induces profibrotic marker expression, consistent with activation of a myofibroblast phenotype.

**Figure 5.**
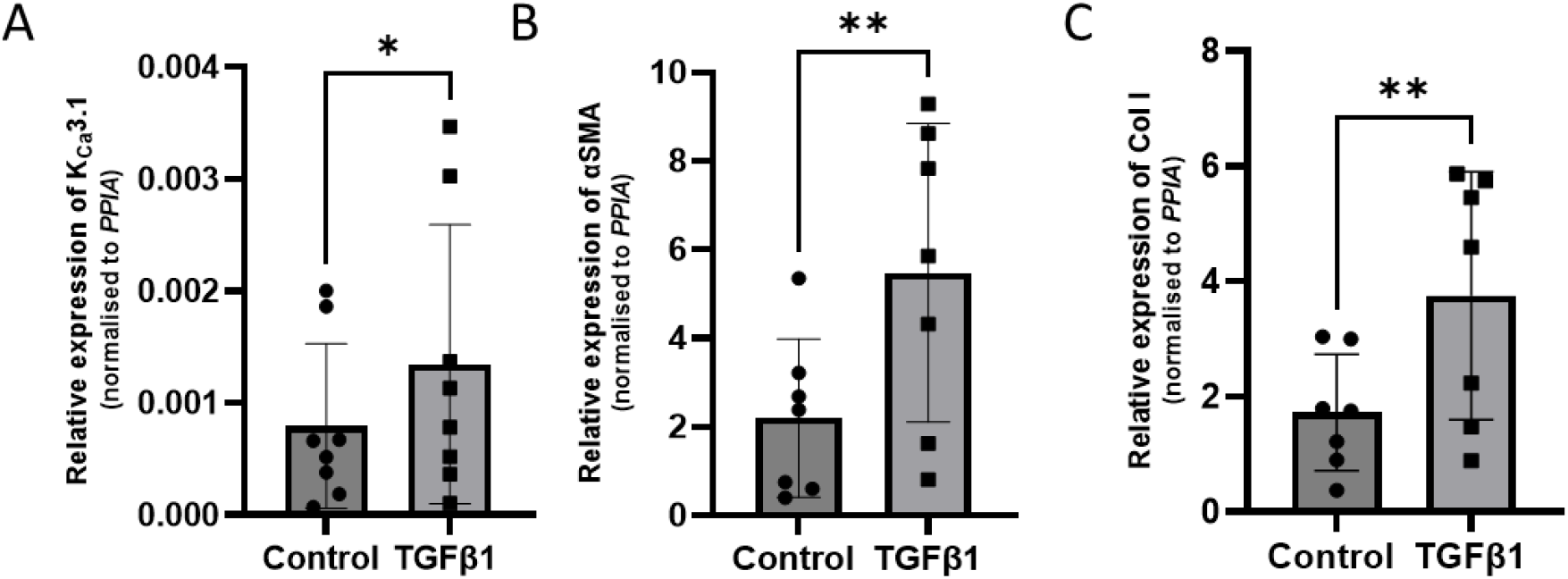
TGFβ1 induces K_Ca_3.1 and profibrotic markers in AV myofibroblasts. qRT-PCR analysis of AV myofibroblasts after 72 h stimulation with TGFβ1 (10 ng/ml), significantly increased mRNA expression of (A) K_Ca_3.1 (*p<0.05, paired t-test), (B) αSMA (**p=0.004, paired t-test) and (C) collagen type I (**p=0.005, paired t-test). Data are normalised to PPIA and presented as mean ± SD, n=8.

### Inhibition of K_Ca_3.1 attenuates the differentiation of AV-derived myofibroblasts

To investigate the role of K_Ca_3.1 in the differentiation of myofibroblasts, we evaluated whether pharmacological blockade of K_Ca_3.1 could modulate TGFβ1-induced myofibroblast activation. Co-treatment with the K_Ca_3.1 inhibitor senicapoc (10 or 100 nM) attenuated the expression of both αSMA and collagen 1 mRNA, with a statistically significant reduction observed for αSMA at 100 nM (p=0.02), whereas the decrease in collagen type I did not reach significance (p=0.09) (**Figure 6A and B**). These results indicate that K_Ca_3.1 contributes to TGFβ1-mediated upregulation of myofibroblast activation and extracellular matrix gene expression.

**Figure 6.**
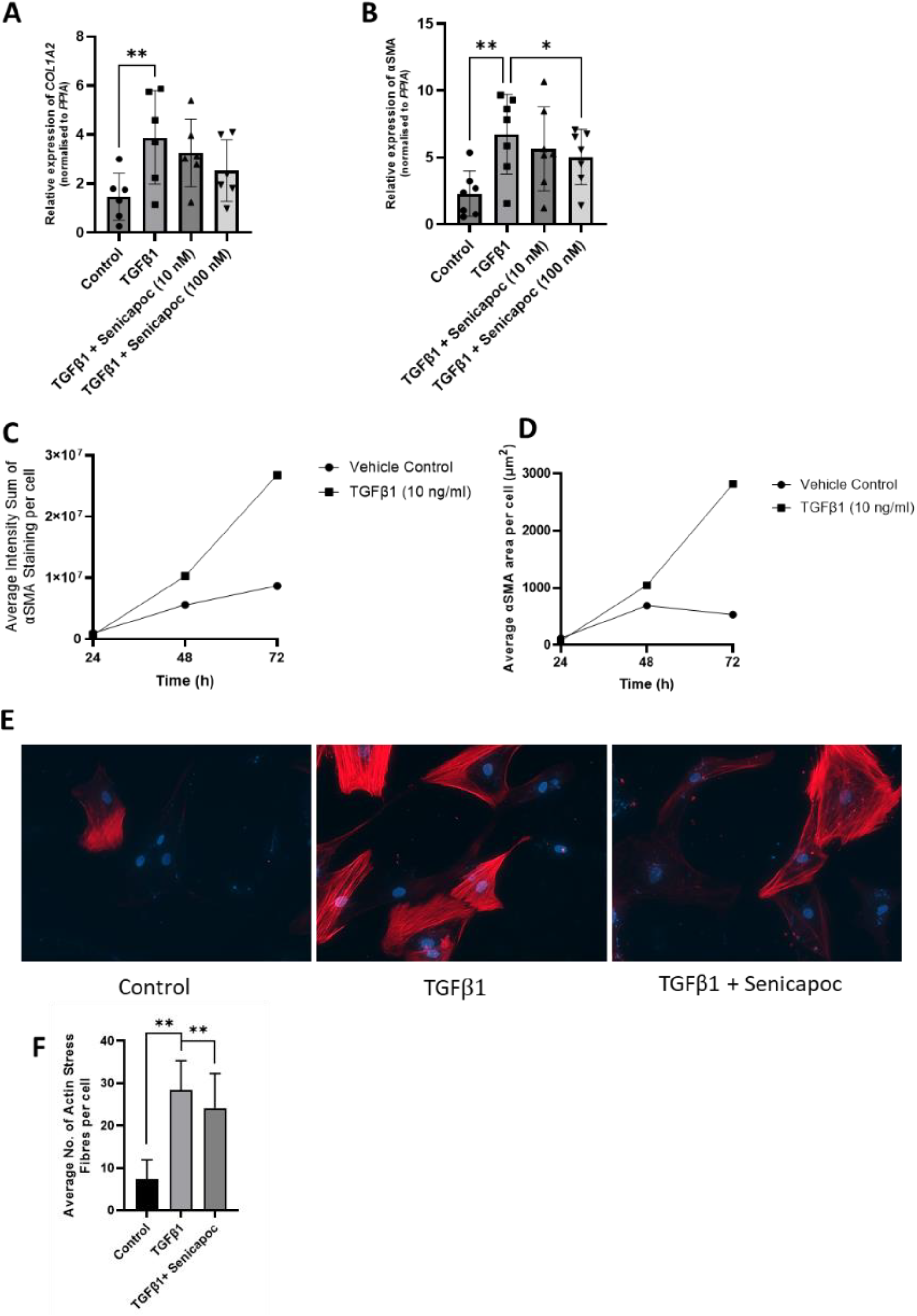
K_Ca_3.1 blockade reduces TGFβ1-induced collagen type I and αSMA expression in AV myofibroblasts. (A) Relative mRNA expression of collagen type I. TGFβ1 significantly increased collagen type I mRNA expression (p=0.005); senicapoc reduced this effect, although not significantly. Gene expression was normalised to PPIA. Data are presented as mean ± SD (n=7). B) Relative mRNA expression for αSMA following 72-hour stimulation with TGFβ1 (10 ng/ml), with or without senicapoc (10 or 100 nM). TGFβ1 significantly increased aSMA mRNA expression (**p=0.004), and co-treatment with 100 nM senicapoc significantly reduced this response (*p=0.02) (n=7). C) Average intensity sum of αSMA staining per cell following 72-hour stimulation with TGFβ1 (10 ng/ml). (D) Average area of αSMA per cell following 72-hour stimulation with TGFβ1 (10 ng/ml). (E) Representative image of AS valve myofibroblasts stained for αSMA (red) and nuclei (DAPI, blue) used for stress fibre analysis. (F) Average number of αSMA stress fibres per cell after 72-hour stimulation with TGFβ1 ± *senicapoc (100 nM). TGFβ1 significantly increased stress fibre formation* (**p=0.005), which was significantly inhibited by senicapoc (**p=0.008, n = 6). Data are shown as mean ± SD. Paired t-test used for statistical analysis.

Time-course experiments revealed that exposure of myofibroblasts to TGFβ1 for 24, 48, and 72 h elicited progressively elevated αSMA expression and enlarged cellular morphology, indicative of sustained myofibroblast activation (**Figure 6C and D**). Stress fibre formation was quantified using an ImageJ-based macro. TGFβ1 stimulation significantly increased αSMA-positive stress fibre formation compared with control (p=0.005, n=6), and this effect was significantly reduced by co-treatment with senicapoc (100 nM, p=0.008) (**Figure 6F**). Representative images are shown in **Figure 6E**. Collectively, these findings suggest that K_Ca_3.1 activity is required for full TGFβ1-induced myofibroblast differentiation and the associated cytoskeletal and ECM remodelling.

### TGFβ1-dependent myofibroblast contraction is reduced by K_Ca_3.1 inhibition

Having demonstrated that K_Ca_3.1 contributes to TGFβ1-induced myofibroblast differentiation, cytoskeletal remodelling, and extracellular matrix gene expression, we next assessed whether K_Ca_3.1 influences the contractile function of AV myofibroblasts. AS valve myofibroblasts were embedded in collagen gels and stimulated with TGFβ1, and gel contraction was monitored over time. Control gels exhibited a time-dependent contraction of 54.7% over 22 hours. TGFβ1 significantly enhanced gel contraction to 77.4% at the final time point. Co-treatment with the K_Ca_3.1 inhibitor senicapoc (100 nM) attenuated TGFβ1-induced contraction, resulting in 66.7% contraction (**Figure 7A**). Quantitative analysis at 22 hours confirmed that TGFβ1 markedly increased contraction relative to controls (*p* <0.0001), and this effect was significantly reduced by senicapoc co-treatment (*p*=0.02) (n=8, **Figure 7B and C**). These findings indicate that K_Ca_3.1 activity contributes not only to the molecular and cytoskeletal hallmarks of myofibroblast differentiation but also to functional contractile responses. By modulating myofibroblast-mediated matrix contraction, K_Ca_3.1 may play a critical role in tissue stiffening and fibrotic remodelling in aortic stenosis.

**Figure 7.**
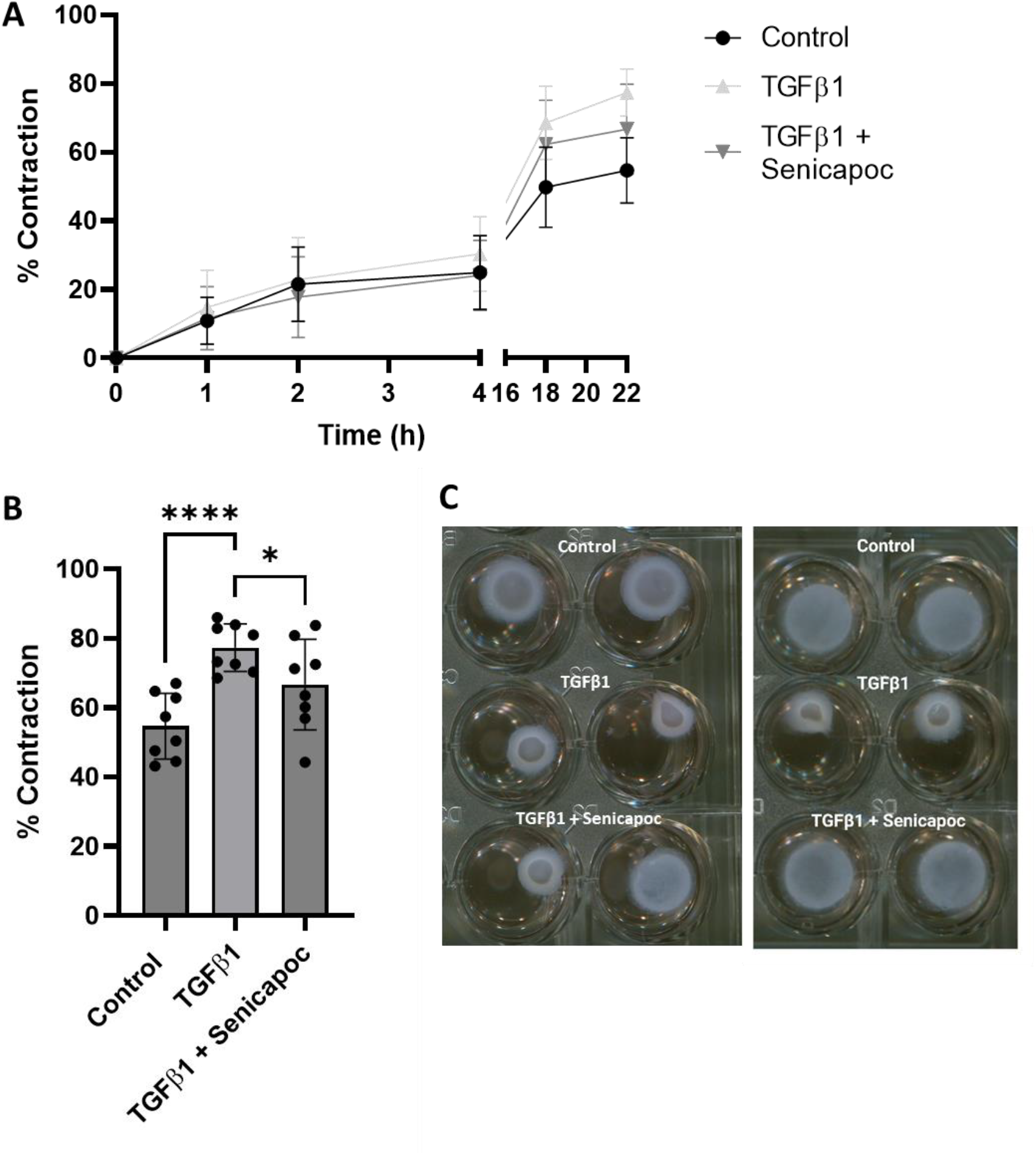
K_Ca_3.1 blockade significantly reduces TGFβ1-induced myofibroblast-mediated contraction. (A) Time course of collagen gel contraction over 22 hours. AS valve myofibroblasts were pre-treated for 24 hours with vehicle (DMSO) or senicapoc (100 nM), then embedded in collagen gels and stimulated with TGFβ1 (10 ng/ml) or vehicle (citric acid). TGFβ1 significantly enhanced gel contraction compared to control, while co-treatment with senicapoc partially reduced this effect. Data represent mean ± SD (n=8). (B) Quantification of gel contraction at 22 hours. TGFβ1 significantly increased contraction compared to control (****p<0.0001), and senicapoc significantly reduced TGFβ1-induced contraction (*P=0.02). Data are shown as mean ± SD (n=8). Paired t-tests were used for statistical analysis. (C) Representative photographs of collagen gels containing AS valve myofibroblasts from two different donors after 22 hours of incubation under the indicated conditions.

### Senicapoc does not inhibit TGFβ1-induced nuclear translocation of Smad2/3

Similar to findings in lung fibrosis[9, 13, 14], we sought to determine which signalling pathways mediate TGFβ1-induced responses in human AV myofibroblasts and whether K_Ca_3.1 contributes to these pathways. In particular, we investigated whether TGFβ1 stimulates the canonical Smad2/3 pathway, a central mediator of profibrotic gene transcription.

Serum-starved AV myofibroblasts were treated with TGFβ1 (10 ng/ml) in the presence or absence of extracellular Ca^2+^. Representative immunofluorescence images (**Figure 8A**) showed a marked increase in nuclear accumulation of Smad2/3 following TGFβ1 stimulation. Quantitative analysis of nuclear-to-whole-cell fluorescence (**Figure 8B**) confirmed a significant increase compared with serum-free controls (p<0.05). Importantly, TGFβ1 stimulation under Ca^2+^-free conditions did not induce Smad2/3 nuclear translocation compared to TGFβ1 treatment in Ca^2+^-containing media (p<0.01), indicating that TGFβ1-mediated nuclear translocation of Smad2/3 is Ca^2+^-dependent. These data indicate that extracellular Ca^2+^ is required for efficient TGFβ1-induced Smad2/3 signalling in human AS valve myofibroblasts.

**Figure 8.**
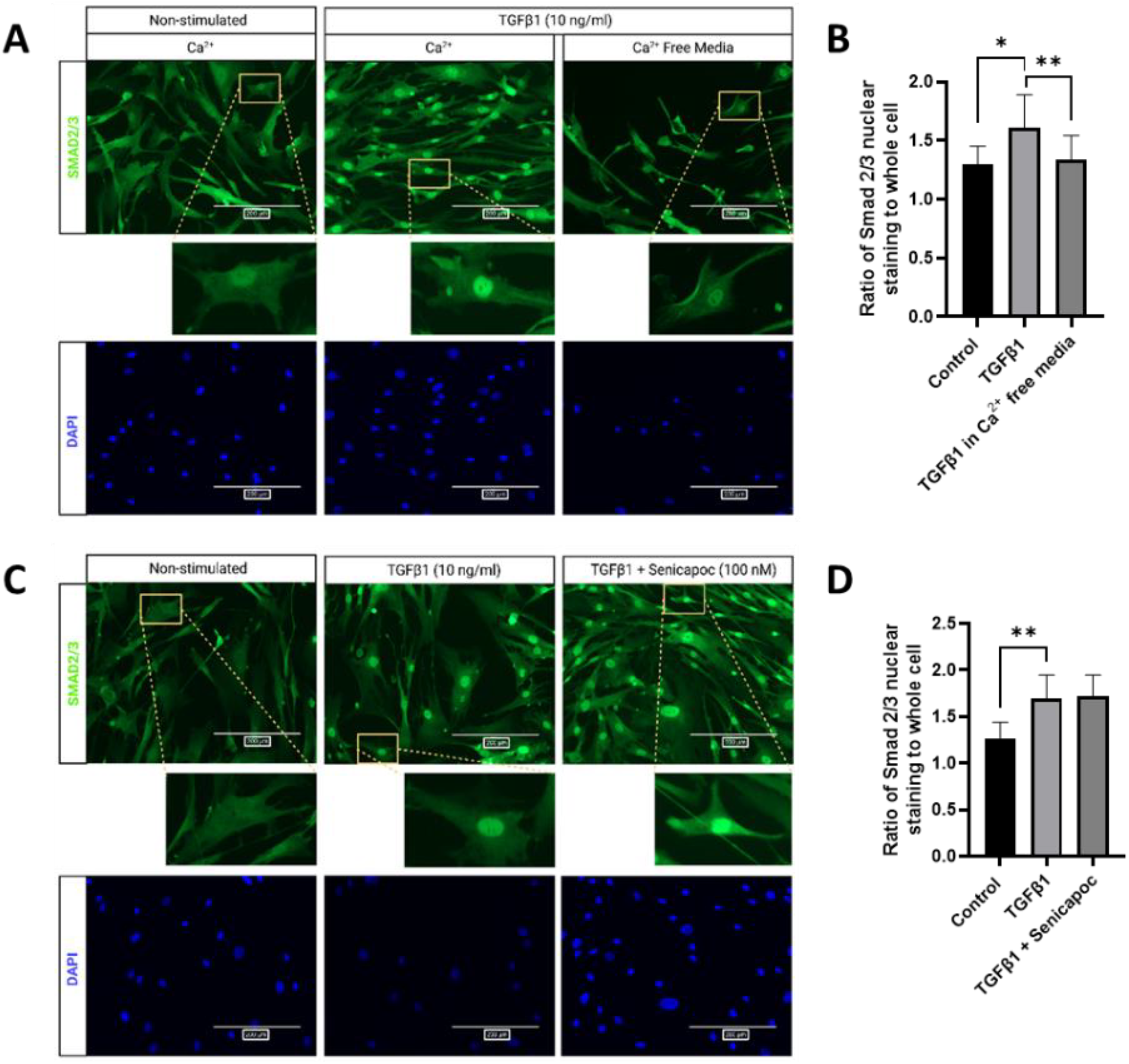
Senicapoc does not inhibit TGFβ1-induced nuclear translocation of Smad2/3 in human AV myofibroblasts. (A) Representative immunofluorescence images of human AV myofibroblasts stained for Smad2/3 (green) and DAPI (blue) under the indicated conditions: non-stimulated, TGFβ1 (10 ng/ml) in Ca^2+^-containing media, and TGFβ1 (10 ng/ml) in Ca^2+^-free media. Insets show higher magnification of individual cells, highlighting nuclear localisation of Smad2/3. Scale bars = 200 μm. (B) Quantification of the ratio of nuclear to whole-cell Smad2/3 fluorescence intensity under the same conditions. TGFβ1 stimulation significantly increased nuclear localisation of Smad2/3 compared to serum-free controls (*p< 0.05). This increase did not occur in the absence of extracellular Ca^2+^ (**p< 0.01). Data represent mean ± SD from 35 cells per condition. (C) Representative immunofluorescence images of human AS valve myofibroblasts stained for Smad2/3 (green) and nuclei with DAPI (blue) under non-stimulated, TGFβ1 stimulation (10 ng/ml), and TGFβ1 stimulation in the presence of senicapoc (100 nM). Insets show magnified views of individual cells, highlighting nuclear localisation of Smad2/3. Scale bars = 200 μm. (D) Quantification of the nuclear-to-whole-cell Smad2/3 fluorescence intensity ratio under the indicated conditions. TGFβ1 significantly increased nuclear Smad2/3 localisation compared to serum-free controls (**p < 0.01). Co-treatment with senicapoc did not significantly alter Smad2/3 nuclear translocation. Data represent mean ± SD from 35 cells per condition.

To test whether K_Ca_3.1 contributes to this pathway, cells were co-treated with TGFβ1 (10 ng/ml) and senicapoc (100 nM). Representative images (**Figure 8C**) and quantitative analysis of the nuclear-to-whole-cell fluorescence ratio (**Figure 8D**) demonstrated that TGFβ1-induced nuclear translocation of Smad2/3 remained unchanged in the presence of senicapoc. These findings suggest that K_Ca_3.1 modulates TGFβ1 signalling via Smad-independent mechanisms.

## Discussion

This study identifies for the first time, the expression of the Ca^2+^-activated K^+^ channel K_Ca_3.1 in both human AS AV leaflets and myofibroblasts derived from this tissue. Pharmacological blockade of K_Ca_3.1 attenuated myofibroblast pro-fibrotic responses, including TGFβ1-induced collagen and αSMA expression, stress fibre assembly and contractility. These findings underscore the contribution of ion channels and electrical activity to valvular remodelling and position K_Ca_3.1 as a potential therapeutic target in AS.

Immunohistochemical analysis demonstrated increased K_Ca_3.1 expression in stenotic aortic valves compared with non-diseased controls, paralleling the increased expression of αSMA, a well-established marker of myofibroblasts. Previous studies demonstrated that αSMA-positive interstitial cells are markedly increased in stenotic valves and are closely associated with fibrotic remodelling and disease progression[15, 16]. Consistent with these observations, we found that K_Ca_3.1 co-localised with αSMA-positive interstitial cells, identifying K_Ca_3.1 as a previously unrecognised feature of activated valve myofibroblasts and implicating this channel in pathological valve remodelling.

To investigate the cellular mechanisms underlying these effects, we established a robust method for isolating primary human valve myofibroblasts directly from stenotic valve tissue. Our approach for isolating human AV myofibroblasts was inexpensive, simple to perform and avoided enzymatic digestion, providing a practical tool for studying these cells. These cells represent the principal effector population responsible for ECM remodelling and leaflet stiffening in AS, providing a physiologically relevant platform to investigate fibrotic mechanisms. Using complementary molecular, imaging, and electrophysiological approaches, we demonstrate that K_Ca_3.1 is expressed and functionally active in AS valve myofibroblasts. Expression varied between donors, mirroring the heterogeneity observed in AV tissue in AS, and potentially differences in disease severity or activation state, consistent with other fibrotic settings such as idiopathic pulmonary fibrosis[9, 17], renal fibrosis[18] and atrial fibrosis[10].

TGFβ1 is a central driver of fibrotic remodelling in AS, promoting myofibroblast differentiation, ECM deposition, and contributing to calcification[5, 19, 20]. We observed that TGFβ1 stimulation increased K_Ca_3.1 expression alongside αSMA and collagen type I, suggesting that K_Ca_3.1 is integrated into profibrotic signalling pathways similar to myofibroblasts in other organs[9, 13, 21-23]. Collagen type I comprises ~70% of the AV ECM[24], while αSMA enhances contractility and tissue stiffening[25]. Mechanistically, K_Ca_3.1 channels regulate membrane potential and Ca^2+^ influx, which are critical determinants of cytoskeletal organisation and contractile function[6]. Inhibition of K_Ca_3.1 significantly reduced TGFβ1-induced αSMA expression, stress fibre formation, and contractility—hallmarks of myofibroblast activation. These findings support a model in which K_Ca_3.1 facilitates Ca^2+^-dependent cytoskeletal remodelling and contractile force generation, processes that directly contribute to leaflet stiffening and disease progression. Consistent with our findings here, inhibition of K_Ca_3.1 with TRAM-34 decreased αSMA expression in myofibroblast-like rat mesangial cells and human lung myofibroblasts[26]. Given that AV myofibroblast contraction contributes significantly to leaflet stiffness[27], K_Ca_3.1 inhibition may mitigate tissue stiffening.

Unlike human lung myofibroblasts, K_Ca_3.1 inhibition did not significantly alter AV myofibroblast proliferation, migration or Smad2/3 nuclear translocation, suggesting that K_Ca_3.1 selectively regulates specific downstream pathways. This contrasts with reports in other fibrotic contexts, such as the human lung, where TRAM-34 reduced lung myofibroblast proliferation following FBS stimulation[9]. Similarly, although K_Ca_3.1 has been linked to Smad signalling in human lung myofibroblasts[13, 14, 28], we observed no effect on Smad2/3 nuclear translocation here. Based on the observed effects on cytoskeletal organisation and contractility, it is likely that K_Ca_3.1 acts through Ca^2+^-sensitive regulators of actin dynamics, such as the Rho/ROCK signalling pathway, which plays a central role in myofibroblast contractile activation[27, 29-31]. This mechanism is consistent with observations in other fibrotic tissues and suggests that K_Ca_3.1 contributes primarily to the mechanical and contractile components of fibrotic remodelling in the AV rather than proliferative responses.

These findings have important translational implications, as there are currently no approved pharmacological therapies capable of slowing the progression of AS. Current management is limited to valve replacement[32]. The identification of K_Ca_3.1 as a regulator of myofibroblast activation and contractile function suggests that pharmacological inhibition of this channel may represent a novel strategy to attenuate fibrotic remodelling and delay disease progression. Senicapoc, a selective K_Ca_3.1 inhibitor, has already undergone extensive clinical evaluation and demonstrated an excellent safety profile, including a 12-month phase III trial in sickle cell disease in which it was well tolerated without major safety concerns[31]. This clinical safety profile substantially lowers the translational barrier for repurposing senicapoc or related K_Ca_3.1 inhibitors in fibrotic cardiovascular disease, and our data thus provide a strong rationale for future translational studies evaluating K_Ca_3.1 as a therapeutic target in AS.

Relevant clinical outcome measures may include emerging imaging biomarkers including CT-derived fibrocalcific scores and valve extracellular matrix remodelling indices as well as AV velocities/gradients[33]. These quantitative imaging tools provide sensitive measures of disease progression and could serve as surrogate endpoints in early-phase clinical trials targeting valvular fibrosis with senicapoc.

This study has several important strengths. First, it provides the first direct demonstration of K_Ca_3.1 expression and functional activity in human AS valve tissue and valve-derived myofibroblasts, using complementary molecular, imaging, and electrophysiological approaches. Second, the use of primary human cells isolated directly from diseased valve tissue enhances the physiological relevance of these findings compared with animal models or immortalised cell lines. Third, functional experiments using pharmacological modulation of K_Ca_3.1 establish a mechanistic role for this channel in regulating myofibroblast differentiation, cytoskeletal organisation, and contractile behaviour. Finally, the use of a clinically validated inhibitor with an established safety profile strengthens the translational relevance of these findings.

Several limitations should be acknowledged. First, this study was performed in vitro using isolated valve myofibroblasts, and therefore does not fully recapitulate the complex multicellular and biomechanical environment of the intact valve. Second, variability in K_Ca_3.1 expression and functional responses between donors likely reflects heterogeneity in disease stage and patient characteristics, although this variability also highlights the relevance of these findings to human disease. Third, while pharmacological inhibition of K_Ca_3.1 attenuated key profibrotic responses, it did not significantly affect all endpoints, such as collagen expression in all donors or proliferative responses, suggesting that additional signalling pathways contribute to valve fibrosis. Finally, the present study did not assess the effects of K_Ca_3.1 inhibition in vivo. Further preclinical and clinical studies are required to determine whether targeting K_Ca_3.1 can alter disease progression in AS.

In conclusion, this study establishes for the first time that K_Ca_3.1 channels are expressed in human AS valve tissue and are functionally active in AS valve-derived myofibroblasts, promoting pro-fibrotic activity. Our findings demonstrate that K_Ca_3.1 activity contributes to valvular fibrosis by regulating TGFβ1-mediated myofibroblast differentiation, cytoskeletal remodelling, and contractile function. These results identify K_Ca_3.1 as a novel regulator of valve remodelling and a promising therapeutic target in AS. Importantly, given the demonstrated clinical safety of senicapoc, these findings provide a strong rationale for future translational and clinical studies evaluating K_Ca_3.1 inhibition as a strategy to slow disease progression and delay the need for valve replacement.

## Author Contributions

Conceived the project: KMR, AS, PB, GM. Obtained ethics approval and set up valve collection: AS, SS. Valve and data extraction/collection: MW, SA, GM, HW, MA, SS, SV, NC. Designed the experiments: KMR, PB, MW. Performed the experiments: MW, KMR, JS, MB, DT, CM, MR, MD, GE. Interpreted the data and wrote and edited the manuscript: KMR, MW, AS, PB, GM. All authors reviewed and edited the manuscript critically for important intellectual content; and gave final approval of the version to be published.

## Acknowledgements

This work was supported by the National Institute for Health and Care Research (NIHR) Leicester Biomedical Research Centre and the Leicester British Heart Foundation Centre of Research Excellence. The views expressed are those of the author(s) and not necessarily those of the NIHR or the Department of Health and Social Care. We gratefully acknowledge Alfie Hall, Jingye Zhou, Keyan Li, and Nikita Kalidas for performing portions of the

experimental work. Imaging support was provided by the Advanced Imaging Facility at the University of Leicester (RRID:SCR_020967). We also thank the cardiac theatre team at Glenfield Hospital for their invaluable assistance with human valve collection.

## Notes

### Competing Interest Statement

The authors have declared no competing interest.

## References

1. Farrar, E.J., et al., Valve interstitial cell tensional homeostasis directs calcification and extracellular matrix remodeling processes via RhoA signaling. Biomaterials, 2016. 105: p. 25–37.

2. Otto, C.M. and B. Prendergast, Aortic-valve stenosis--from patients at risk to severe valve obstruction. N Engl J Med, 2014. 371(8): p. 744–56.

3. Ji, H., et al., Rho/Rock cross-talks with transforming growth factor-β/Smad pathway participates in lung fibroblast-myofibroblast differentiation. Biomed Rep, 2014. 2(6): p. 787–792.

4. Huang, C., et al., Blockade of KCa3.1 ameliorates renal fibrosis through the TGF-beta1/Smad pathway in diabetic mice. Diabetes, 2013. 62(8): p. 2923–2934.

5. Varshney, R., et al., Inactivation of platelet-derived TGF-β1 attenuates aortic stenosis progression in a robust murine model. Blood Adv, 2019. 3(5): p. 777–788.

6. Roach, K.M. and P. Bradding, Ca(2+) signalling in fibroblasts and the therapeutic potential of KCa 3.1 channel blockers in fibrotic diseases. Br J Pharmacol, 2019.

7. Simard, C., et al., Ion Channels in the Development and Remodeling of the Aortic Valve. Int J Mol Sci, 2023. 24(6).

8. Hamill, O.P., et al., Improved patch-clamp techniques for high-resolution current recording from cells and cell-free membrane patches. Pflugers Archiv : European journal of physiology, 1981. 391(2): p. 85–100.

9. Roach, K.M., et al., The K+ Channel KCa3.1 as a Novel Target for Idiopathic Pulmonary Fibrosis. PLoS ONE, 2013. 8(12): p. e85244.

10. Wang, Z., et al., Elevated K(Ca)3.1 expression by angiotensin II via the ERK/NF-κB pathway contributes to atrial fibrosis. J Mol Cell Cardiol, 2025. 202: p. 133–143.

11. Zheng, K.H., E. Tzolos, and M.R. Dweck, Pathophysiology of Aortic Stenosis and Future Perspectives for Medical Therapy. Cardiol Clin, 2020. 38(1): p. 1–12.

12. Ataga, K.I., et al., Efficacy and safety of the Gardos channel blocker, senicapoc (ICA-17043), in patients with sickle cell anemia. Blood, 2008. 111(8): p. 3991–3997.

13. Roach, K.M., et al., Human lung myofibroblast TGFbeta1-dependent Smad2/3 signalling is Ca(2+)-dependent and regulated by KCa3.1 K(+) channels. Fibrogenesis & tissue repair, 2015. 8: p. 5–015-0022-0. eCollection 2015.

14. Roach, K.M., et al., Increased constitutive αSMA and Smad2/3 expression in idiopathic pulmonary fibrosis myofibroblasts is KCa3.1-dependent. Respir.Res., 2014. 15(155): p. doi:10.1186/s12931-014-0155-5.

15. Patterson, A.J.S., et al., Extracellular Matrix Dynamics in Aortic Valve Health and Disease: Insights into Fibrocalcific Remodeling and Creation of Biomimetic Platforms. J Heart Valve Soc, 2024. 1(1).

16. Rabkin-Aikawa, E., et al., Dynamic and reversible changes of interstitial cell phenotype during remodeling of cardiac valves. J Heart Valve Dis, 2004. 13(5): p. 841–7.

17. Organ, L., et al., Inhibition of the K(Ca)3.1 Channel Alleviates Established Pulmonary Fibrosis in a Large Animal Model. American Journal of Respiratory Cell and Molecular Biology, 2017. 56(4): p. 539–550.

18. Grgic, I., et al., Renal fibrosis is attenuated by targeted disruption of KCa3.1 potassium channels. Proceedings of the National Academy of Sciences of the United States of America, 2009. 106(34): p. 14518–14523.

19. Jian, B., et al., Progression of aortic valve stenosis: TGF-β1 is present in calcified aortic valve cusps and promotes aortic valve interstitial cell calcification via apoptosis. The Annals of Thoracic Surgery, 2003. 75(2): p. 457–465.

20. Sritharen, Y., et al., Pathophysiology of Aortic Valve Stenosis: Is It Both Fibrocalcific and Sex Specific? Physiology (Bethesda), 2017. 32(3): p. 182–196.

21. Huang, C., et al., KCa3.1 mediates activation of fibroblasts in diabetic renal interstitial fibrosis. Nephrology Dialysis Transplantation, 2014. 29(2): p. 313–324.

22. Ju, C.-H., et al., Blockade of KCa3.1 Attenuates Left Ventricular Remodeling after Experimental Myocardial Infarction. Cellular Physiology and Biochemistry, 2015. 36(4): p. 1305–1315.

23. Freise, C., et al., K+-channel inhibition reduces portal perfusion pressure in fibrotic rats and fibrosis associated characteristics of hepatic stellate cells. Liver International, 2015. 35(4): p. 1244–1252.

24. Eriksen, H.A., et al., Type I and type III collagen synthesis and composition in the valve matrix in aortic valve stenosis. Atherosclerosis, 2006. 189(1): p. 91–8.

25. David Merryman, W., et al., The effects of cellular contraction on aortic valve leaflet flexural stiffness. Journal of Biomechanics, 2006. 39(1): p. 88–96.

26. Simard, C., et al. Ion Channels in the Development and Remodeling of the Aortic Valve. International Journal of Molecular Sciences, 2023. 24, 5860 DOI: 10.3390/ijms24065860.

27. Jian, B., et al., Progression of aortic valve stenosis: TGF-beta1 is present in calcified aortic valve cusps and promotes aortic valve interstitial cell calcification via apoptosis. Ann Thorac Surg, 2003. 75(2): p. 457–65; discussion 465-6.

28. Yu, Z., et al., Targeted inhibition of KCa3.1 attenuates TGF-beta-induced reactive astrogliosis through the Smad2/3 signaling pathway. Journal of neurochemistry, 2014. 130(1): p. 41–49.

29. Fu, R.G., et al., Inhibition of the K+ channel K(Ca)3.1 reduces TGF-β1-induced premature senescence, myofibroblast phenotype transition and proliferation of mesangial cells. PLoS One, 2014. 9(1): p. e87410.

30. Morotti, M., et al., Muscle Damage in Dystrophic mdx Mice Is Influenced by the Activity of Ca(2+)-Activated K(Ca)3.1 Channels. Life (Basel), 2022. 12(4).

31. Osman, N., et al., Smad2-dependent glycosaminoglycan elongation in aortic valve interstitial cells enhances binding of LDL to proteoglycans. Cardiovasc Pathol, 2013. 22(2): p. 146–55.

32. Strange, G.A., et al., Uncovering the treatable burden of severe aortic stenosis in the UK. Open Heart, 2022. 9(1).

33. Lembo, M., et al., Quantitative Computed Tomography Angiography for the Evaluation of Valvular Fibrocalcific Volume in Aortic Stenosis. JACC Cardiovasc Imaging, 2024. 17(11): p. 1351–1362.

